# Ex vivo Demonstration of Functional Deficiencies in Popliteal Lymphatic Vessels from TNF-Tg Mice with Inflammatory Arthritis

**DOI:** 10.1101/2020.09.22.309070

**Authors:** Joshua P. Scallan, Echoe M. Bouta, Homaira Rahimi, H. Mark Kenney, Christopher T. Ritchlin, Michael J. Davis, Edward M. Schwarz

## Abstract

**Background:** Rheumatoid arthritis (RA) is a progressive immune-mediated inflammatory disease characterized by intermittent episodes of pain and inflammation in affected joints, or flares. Recent studies demonstrated lymphangiogenesis and expansion of draining lymph nodes during chronic inflammatory arthritis, and lymphatic dysfunction associated with collapse of draining lymph nodes in RA patients and TNF-transgenic (TNF-Tg) mice experiencing arthritic flare. As the intrinsic differences between lymphatic vessels afferent to healthy, expanding, and collapsed draining lymph nodes are unknown, we characterized the ex vivo behavior of popliteal lymphatic vessels (PLVs) from WT and TNF-Tg mice. We also interrogated the mechanisms of lymphatic dysfunction through inhibition of nitric oxide synthase (NOS).

**Methods:** Popliteal lymph nodes (PLNs) in TNF-Tg mice were phenotyped as Expanding or Collapsed by in vivo ultrasound and age-matched to WT littermate controls. The PLVs were harvested and cannulated for ex vivo functional analysis over a relatively wide range of hydrostatic pressures (0.5 to 10 cmH_2_O) to quantify the end diastolic diameter (EDD), tone, amplitude (AMP), ejection fraction (EF), contraction frequency (FREQ) and fractional pump flow (FPF) with or without NOS inhibitors Data was analyzed using repeated measures two-way ANOVA with Bonferroni’s *post hoc* test.

**Results:** Real time videos of the cannulated PLVs demonstrated the predicted phenotypes of robust versus weak contractions of the WT versus TNF-Tg PLV, respectively. Quantitative analyses confirmed that TNF-Tg PLVs had significantly decreased AMP, EF and FPF versus WT (p<0.05). EF and FPF were recovered by NOS inhibition, while the reduction in AMP was NOS independent. No differences in EDD, tone, or FREQ were observed between WT and TNF-Tg PLVs, nor between Expanding versus Collapsed PLVs.

**Conclusion:** These findings support the concept that chronic inflammatory arthritis leads to NOS dependent and independent draining lymphatic vessel dysfunction that exacerbates disease, and may trigger arthritic flare due to decreased egress of inflammatory cells and soluble factors from affected joints.

## INTRODUCTION

Rheumatoid arthritis (RA) is an inflammatory joint disease that affects 0.5-1% of the population (1, 2). While there have been major advances in our understanding of RA pathogenesis, significant unmet clinical needs remain for many patients who are refractory to available treatments (3, 4). Although specific autoantibodies are diagnostic of RA, it is now broadly accepted that environmental and epigenetic factors are also critical in the development and pathogenesis of joint disease (5). Additionally, the mechanisms of inflammatory joint disease in RA patients who do not have detectable levels of these autoantibodies (seronegative RA) are poorly understood (6). Thus, research is needed to elucidate non-autoimmune etiologies of RA.

Another major challenge for RA patients and caregivers is the tendency of the disease to flare, and the relative refractory nature of persistent disease despite aggressive therapy (5). Although autoimmune mechanisms underlie the development of RA, the pathways that trigger flare are not well understood, and alternative pathways may contribute to disease exacerbation (7). Although recent gene expression studies have identified potential roles of immature IgD^+^ B-cells, circulating CD45^−^/CD31^−^/PDPN^+^ pre-inflammatory mesenchymal (PRIME) cells, and synovial macrophages in RA flare (8, 9), it is also possible that exacerbation of disease is due to the loss of protective mechanisms in the setting of chronic joint inflammation. One area of interest receiving increased attention in RA is the synovial lymphatic system and its dysfunction, based on animal models and clinical pilots, where the loss of lymphatic drainage of inflamed joints is strongly implicated in the onset of synovitis and disease progression (7, 10, 11).

Previously, we demonstrated that arthritic progression in knee joints of tumor necrosis factor-transgenic (TNF-Tg) mice is paralleled by dramatic changes in the draining lymph nodes (12-14). These longitudinal imaging studies combining contrast enhanced (CE) MRI of the synovium and popliteal lymph node (PLN) (12), with quantification of lymphatic drainage via near infrared (NIR) imaging of an injected dye (indocyanine green, ICG) (15), demonstrated that prior to detectable synovial hyperplasia in the knee, the adjacent PLN expands. This PLN expansion is associated with increased lymphangiogenesis, elevated volume, CD11b^+^ macrophage infiltration, and the accumulation of a unique subset of IgD^+^/CD23^+^/CD21^hi^ <u>B</u> cells in inflamed <u>n</u>odes (B-in) (15-21). This asymptomatic “expansion” phase is followed by a sudden “collapse” of the PLN, which is objectively defined by quantitative CE-MRI or power Doppler ultrasound (PD-US) imaging of the PLN (12, 22). This collapse, which occurs at variable time intervals in ∼80% of TNF-Tg mice, is associated with B-in translocation from the lymph node follicles to LYVE-1^+^ lymphatic vessels of the paracortical sinuses. Thereafter, lymphatic drainage declines significantly due to loss of intrinsic lymphatic contractions and passive flow (14, 16, 20, 23). It was also demonstrated that B cell depletion therapy (BCDT) with anti-CD20 antibodies ameliorated knee flare by “clearing” the collapsed lymph node sinuses, and restoring passive lymphatic flow despite the continued absence of detectable lymphatic contractions (14).

In addition, we have previously reported that nitric oxide signaling is associated with the loss of the lymphatic pulse in TNF-Tg mice. Pharmacologic inhibition of inducible nitric oxide synthase (iNOS) using L-N^6^-(1-iminoethyl)lysine 5-tetrazole-amide (L-NIL) in TNF-Tg mice has been shown to recover PLV contractions and ICG clearance via NIR imaging (24). We have also recently demonstrated that PLN expansion during arthritic progression is dependent on iNOS by longitudinally monitoring PLN volumes with PD-US in TNF-Tg versus TNF-Tg x iNOS^−/−^ mice (25). Global ablation of iNOS preserved PLV contraction frequency assessed by in vivo NIR-ICG imaging, which was associated with significantly reduced synovitis in female TNF-Tg mice that typically exhibit an accelerated disease course (25, 26). Thus, the specific features associated with nitric oxide signaling and PLV contractility in TNF-Tg mice warrants further investigation.

Additionally, lymphatic dysfunction in RA has also been demonstrated in clinical pilot studies. Manzo et al used PD-US to show that joint draining lymph nodes in RA patients are subjected to subclinical intra-parenchymal changes and vascular flow modulation (27). Subsequently, we used CE-MRI to show that RA patients with the smallest change in lymph node volume during anti-TNF therapy experienced the greatest pain relief in symptomatic knee joints with a remarkably linear inverse correlation (28). Most recently, we utilized NIR-ICG imaging (10) to show that lymphatic drainage in the hands of RA patients with active disease is decreased compared to healthy controls (29). Furthermore, this reduced lymphatic drainage was associated with a decrease in the total length of ICG^+^ lymphatic vessels on the dorsal surface of the hands, which continued to contract at a similar rate as controls (29). Collectively, these findings support the hypothesis that there is an intrinsic defect in RA joint draining lymphatic vessels in advanced disease and/or flare, which warrants direct testing via ex vivo functional analyses. To this end, we utilized our ex vivo approaches to assess the contractile function of PLVs from WT and TNF-Tg mice under conditions of controlled hydrostatic and oncotic pressures in the absence of flow (30-32). Moreover, real-time diameter tracking of ex vivo PLV pump function in this system allows for quantification of end diastolic diameter (EDD), tone, amplitude (AMP), ejection fraction (EF), contraction frequency (FREQ), and fractional pump flow (FPF). We also assessed alterations in these outcome measures in WT and TNF-Tg PLVs following non-specific NOS inhibition using Nω-nitro-L-arginine methyl ester (L-NAME). Here we report initial findings with this methodology, which demonstrate progressive intrinsic defects in PLVs from TNF-Tg mice with inflammatory-erosive arthritis, some of which are dependent on NOS activity.

## MATERIALS AND METHODS

### Animals and treatment

All animal research was conducted on IACUC approved protocols. TNF-Tg mice (3647 line) (33) were originally acquired from Dr. G. Kollias, and are maintained as heterozygotes in a C57BL/6 background. For all imaging, mice were anesthetized with 1.5-2% isoflurane. For tissue harvesting, mice were anesthetized with pentobarbital sodium and immediately following this procedure, euthanized by overdose.

### Power Doppler ultrasound (PD-US)

The PLNs of TNF-Tg mice were phenotyped as “Expanding” or “Collapsed” using PD-US as previously described (22). PD-US was also performed on the knee joints to confirm disease severity as previously described (34). Each joint was imaged with a high-resolution small-animal ultrasound system (VisualSonics 770, Toronto, Ontario, Canada) using a 704b scanhead.

### Near infrared indocyanine green (NIR-ICG) imaging

Mice were placed on a heated surface (Indus Instruments, Webster, TX, USA), hair was removed with a depilatory cream and the mouse footpad was injected with 10 μl of 0.1% ICG (Akorn, Lake Forest, Illinois, USA) as previously described (35). The imaging system was composed of a lens (Zoom 7000, Navitar, Rochester, NY, USA), ICG filter set (Semrock, Lake Forest, IL, USA) and camera (Prosilica GT1380, Allied Vision Technologies, Exton, PA, USA). ICG was excited with a tungsten halogen bulb (IT 9596ER, Illumination Technologies, Inc., Syracuse, NY, USA) through a ring illuminator (Schott, Elmsford, NY, USA). Imaging settings and recordings were accomplished through a custom built LabVIEW program (National Instruments, Austin, TX, USA). Real time NIR imaging was performed for 60 minutes after ICG injection into the footpad to quantify the lymphatic contraction rate, and mice were imaged 24 hours later to quantify % lymphatic clearance as previously described (35).

### Vessel isolation procedure

PLVs (n=5) were harvested from WT and TNF-Tg mice as previously described (36). Briefly, the mice were anaesthetized with pentobarbital sodium (Nembutal; 60 mg kg^−1^, i.p.) and placed in the prone position on a heating pad. An anterolateral incision (∼1 cm) was made in the skin beginning at the ankle of one leg to expose the two PLVs adjacent to the superficial saphenous vein. After the connective tissue on either side of the vein was cleared away, the more superficial of two PLVs was then separated from the vein and placed in Krebs buffer containing albumin. Afterwards, the animal was euthanized by an overdose of pentobarbital sodium (200 mg kg^−1^, i.p.). PLVs (∼40–80 μm i.d.; 1–2 mm long) were pinned in a Sylgard dish and cleaned of connective and adipose tissue before transfer to a 3 mL chamber where the vessel was cannulated, pressurized, and trimmed of any remaining connective tissue prior to beginning the experimental protocol.

### Solutions and chemicals

Krebs buffer contained (in mm): NaCl, 146.9; KCl, 4.7; CaCl_2_·2H_2_O, 2; MgSO_4_, 1.2; NaH_2_PO_4_·H_2_O, 1.2; NaHCO_3_, 3; sodium-Hepes, 1.5; d-glucose, 5 (pH 7.4 at 37°C). An identical buffer was prepared with the addition of 0.5% BSA. During cannulation, the luminal and abluminal solutions contained Krebs with BSA, but during the experiment the abluminal solution was constantly exchanged with fresh Krebs lacking BSA. For the L-NAME experiments, L-NAME was added to the Krebs buffer lacking BSA at a concentration (1 ⨯ 10^−4^), which is sufficient to maximally inhibit eNOS activity (36-38). At the end of every experiment, a Ca^2+^-free physiological saline solution was used to obtain the passive diameter (39). All chemicals were obtained from Sigma (St Louis, MO, USA), with the exception of BSA (US Biochemicals; Cleveland, OH, USA), MgSO_4_ (Fisher Scientific; Pittsburgh, PA, USA) and sodium-Hepes (Fisher Scientific).

### Pressure control and data acquisition

Vessels were cut to a length that contained only a single valve. To prevent continuous, but not pulsatile, flow through the vessel during the experiment, input and output pressures were kept equal (38, 40). PLV segments were tied onto two glass micropipettes (40 μm o.d.) mounted on a Burg-style V-track system (41).

Polyethylene tubing (PE-190) attached to each micropipette was later connected to a valve that allowed pressure control to be switched between a manual reservoir and servo-controlled pumps (39). After the isolated vessel chamber was positioned on an inverted microscope, a suffusion line connected to a peristaltic pump maintained a constant superfusion of Krebs buffer at a rate of 0.4 ml min^−1^; a second line attached to the peristaltic pump in reverse orientation was used to remove excess buffer at the same rate. Input and output pressures were set briefly to the highest pressure used in this study (10 cmH_2_O) to facilitate the removal of axial slack, which minimized bowing of the vessel at high pressures that otherwise interfered with diameter tracking. Afterward, both pressures were lowered to 3 cmH_2_O to allow the vessel to warm up to 37°C and begin contracting.

The input and output pressure transducer signals were recorded on a computer, and displayed a video image of the vessel using a firewire camera (model A641FM Basler; Ahrensburg, Germany) at 30 Hz. A custom-written LabView program (National Instruments; Austin, TX, USA) measured the inner diameter (i.d.) of the vessel on the video image and recorded it as a function of time (39). Each PLV was equilibrated for 30-60 min at 2–3 cmH_2_O and 37°C until a stable pattern of spontaneous contractions developed. Spontaneous contractions were recorded at each pressure for 2–6 min, a time sufficient to obtain at least three contractions at the lowest pressure of 0.5 cmH_2_O. After the last pressure step to 10 cmH_2_O, pressures were lowered to 3 cmH_2_O and the contraction pattern was allowed approximately 20 min to stabilize. All diameters reported here are inner diameters, and end diastolic diameter (EDD), tone, amplitude (AMP), ejection fraction (EF), contraction frequency (FREQ) and fractional pump flow (FPF) were quantified as previously described (36).

### Statistical analysis

Data obtained from the pressure step protocol were plotted as a function of pressure (cmH_2_O). Raw pressure/diameter traces were plotted against time using Igor Pro (Wavemetrics, Lake Oswego, OR, USA). To compare responses obtained within the same vessel, a repeated-measures two-way ANOVA was used in conjunction with Bonferroni’s *post hoc* test. All data were tabulated using Excel, and statistical tests were performed using Prism 5 (Graphpad Software Inc., CA, USA), with significance for all tests set at *P* < 0.05 and reported as means (±SEM).

## RESULTS

### Ex Vivo Popliteal Lymphatic Vessels from TNF-Tg Mice Exhibit Distinct Functional Deficits

Gross visualization of WT (Movie 1), Expanded (Movie 2) and Collapsed (Movie 3) PLV contractions revealed that the TNF-Tg PLVs appeared to be larger in diameter, and had much weaker/ineffective contractions compared to WT, but no differences in contraction frequency were observed (Figure 1). Although quantification of the end diastolic diameter (EDD) failed to detect significant differences between the groups, it did reveal a trend of increased EDD of Collapsed PLV at all pressures tested (Figure 2A). We also observed a consistent trend of decreased tone in both Expanding and Collapsed PLV versus WT at all pressures, which did not reach statistical significance (Figure 2B). The combination of increased EDD and decreased tone likely counteracted each other leading to data that were not significant.

**Figure 1.**
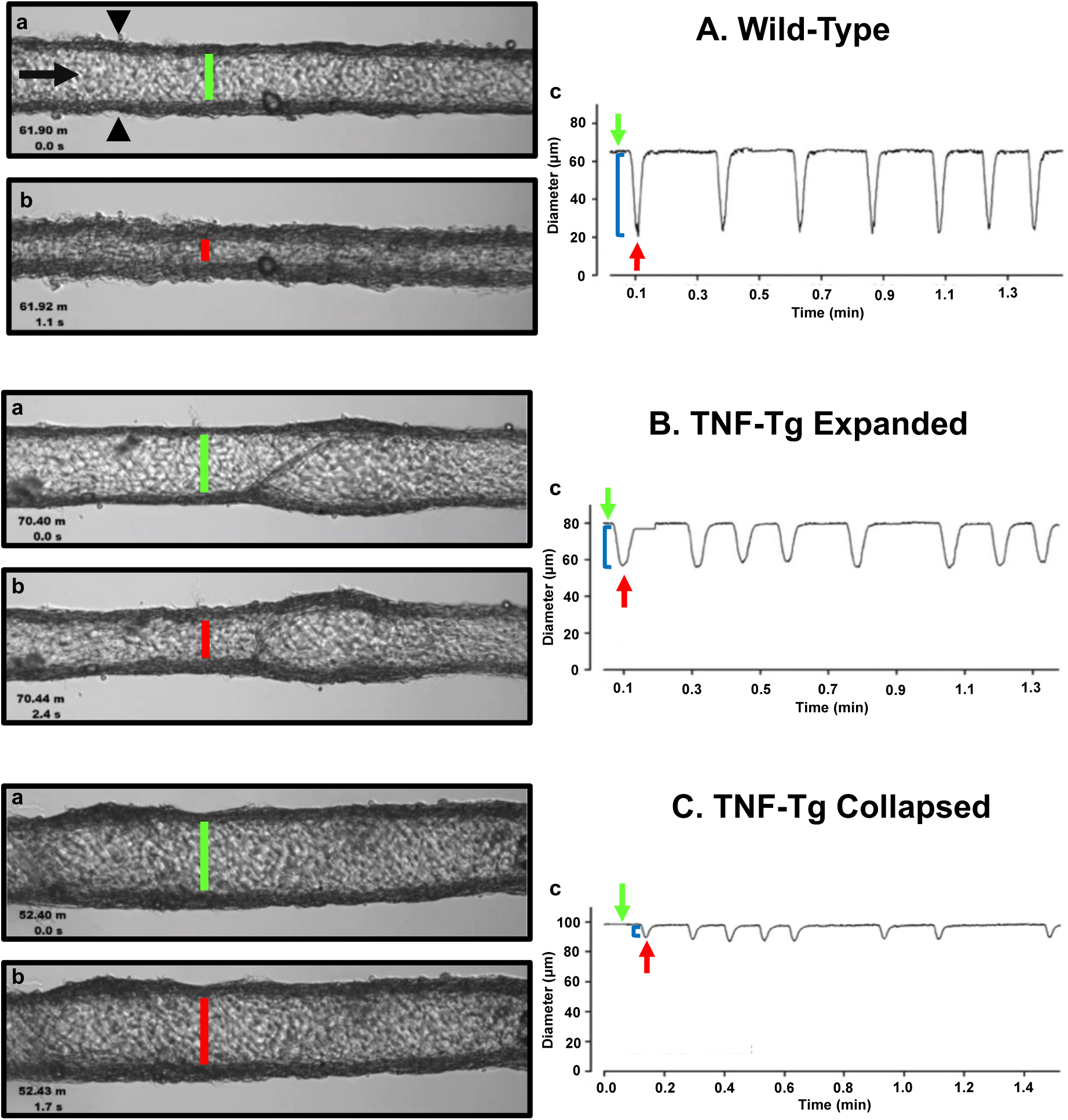
Isolated popliteal lymphatic vessels from TNF-Tg mice exhibit a progressive reduction in contractility. Popliteal lymphatic vessels (PLVs) were harvested from wild-type **(A)** and TNF-Tg mice with expanded **(B)** and collapsed **(C)** popliteal lymph nodes (n = 5 mice per group). The PLVs were then cannulated with constant fluid flow into the lumen of the vessel (black arrow) to measure spontaneous PLV contractions by assessing changes in diameter between the vessel walls (black arrowheads) **(A**.**a)**. In each condition, representative PLVs are shown in diastole **(A-C**.**a)** and systole **(A-C**.**b)** with quantified changes in diameter representing contractions (local minima) over time under constant pressure (3 cmH_2_0) **(A-C**.**c)**. Note the increase in end diastolic diameter (EDD; green lines) and end systolic diameter (ESD; red lines) in TNF-Tg PLVs (expanded < collapsed) compared to wild-type PLVs **(A-C**.**a**,**b)**. The change in EDD (green arrows) and ESD (red arrows) represents a reduction in contraction amplitude (blue brackets) in TNF-Tg PLVs (expanded > collapsed) relative to wild-type control PLVs **(A-C**.**c)**.

Formal quantification of the PLV contractions confirmed the apparent deficiencies in contractile amplitude (AMP) and ejection fraction (EF) observed in the videos. The contraction AMP and EF of both Expanding and Collapsed PLVs were significantly decreased versus WT at pressures of 0.5 to 5.0 cmH_2_O (P<0.05) (Figure 2C and D). Also consistent with the videos, we did not observe differences in PLV contraction frequency at any pressure (Figure 2E). However, despite normal contraction frequency, Expanding and Collapsed PLVs showed significantly decreased FPF at pressures of 2.0 to 5.0 cmH_2_O versus WT (Figure 2F), demonstrating the severity of the TNF-induced defect in PLV contraction strength under these conditions.

**Figure 2.**
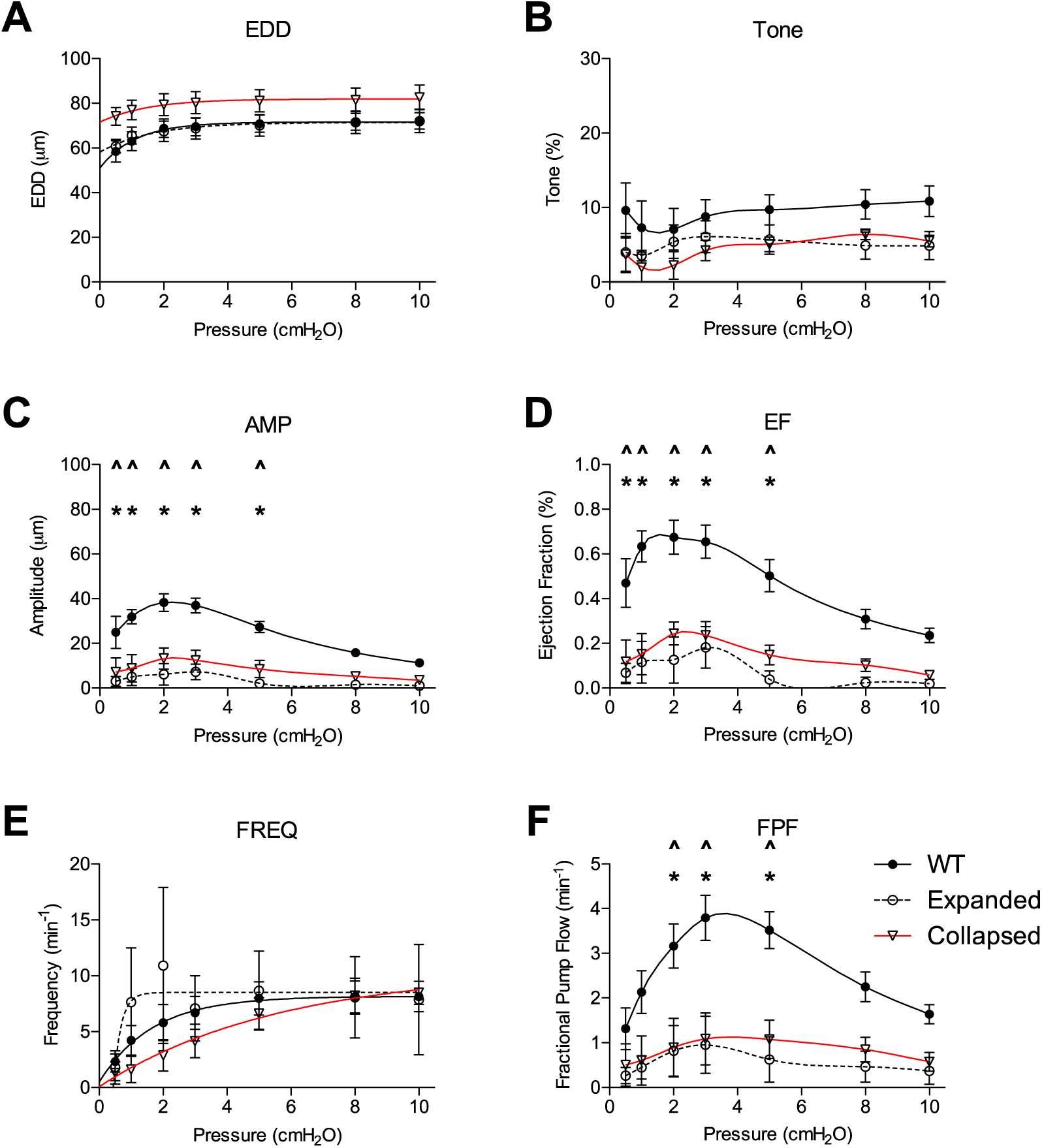
Expanding and Collapsed TNF-Tg mice show reduced amplitude (AMP), ejection fraction (EF), and fractional pump flow (FPF) compared to WT. Lymphatic contractile parameters are plotted against pressure on the *x*-axis, and include end diastolic diameter (EDD; *A*), tone (*B*), contraction amplitude (AMP; *C*), ejection fraction (EF; *D*), contraction frequency (FREQ; *E*) and FPF (*F*). All data are means (± SEM). When error bars appear missing, they are actually contained within the data points. *Filled *versus* open data points differ significantly (*P* < 0.05); ‡filled and open data points both differ from their respective first data point at 0.5 cmH_2_O; †only filled data points differ significantly from the first data point at 0.5 cmH_2_O.

### Functional Defects in TNF-Tg Popliteal Lymphatic Vessels Are Mediated by Nitric Oxide-Dependent Mechanisms

To better understand the functional deficits noted in the TNF-Tg PLVs compared to WT controls, we assessed the role of nitric oxide in mediating the reduced contractility by treating the PLVs with and without L-NAME. In WT and Expanding PLVs, L-NAME administration did not produce any notable changes in tone, while Collapsed PLVs exhibited significantly increased tone at all pressures (except 2.0 cmH_2_O) (Figure 3A – C). Similarly, contraction frequency remained unchanged in WT (except 0.5 cmH_2_O) and Expanding PLVs, whereas Collapsed PLVs showed significantly increased contraction frequency at all pressures after treatment with L-NAME (Figures 3D – F). Following L-NAME treatment, WT and Expanding PLVs demonstrated a significant increase in FPF at pressures ≤ 3.0 cmH_2_O (Figure 3G & H). However, Collapsed PLVs demonstrated significant increases in FPF at all pressures following L-NAME administration (Figures 3I). These results indicate a substantial role for nitric oxide signaling in the reduction of PLV contractility during the Collapsed phase of TNF-mediated inflammatory arthritis.

**Figure 3.**
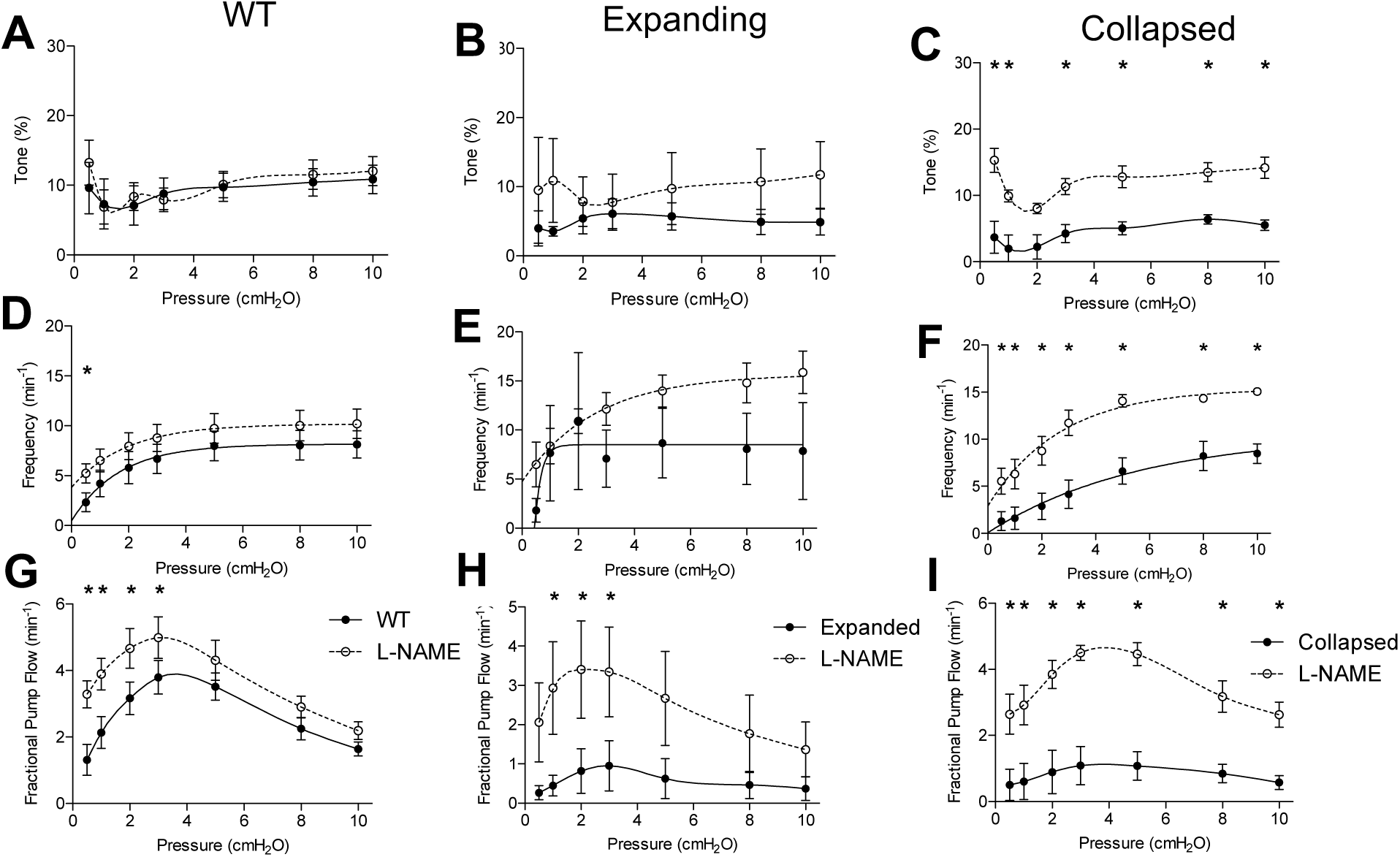
Collapsed lymphatic vessels display distinct nitric oxide-dependent functional defects ex vivo. The lymphatic vessels described in Figure 1 were treated with and without L-NAME (10^−4^ M), and their tone, contraction frequency and fractional pump flow versus pressure were determined ex vivo. All data are means (± SEM). When error bars appear missing, they are actually contained within the data points (\**P* < 0.05 vs. L-NAME at the same pressure). Note that L-NAME has very limited effects on vessels afferent to WT and Expanding PLN, but restores tone, frequency, and fractional pump flow in lymphatic vessels afferent to Collapsed PLN at all pressures tested.

Nitric oxide synthase inhibition improved many of the functional defects in TNF-Tg PLVs when compared to untreated WT levels (Figure 4). As expected from the untreated PLVs (Figure 2), EDD, and thus % tone, remained unchanged between the groups after L-NAME administration (Figure 4A & B). L-NAME also reversed the contractile defects in AMP noted in Collapsed PLVs at pressures of 0.5 – 5.0 cmH_2_O (Figure 2C vs. Figure 4C). However, L-NAME was unable to significantly correct the contractile defect in AMP in Expanding PLVs at the same pressures (Figure 4C), indicating the presence of a nitric oxide-independent mechanism that may also contribute to contractile defects prior to the Collapsed stage. Both EF and FPF, which showed significant reductions in both Expanding and Collapsed TNF-Tg PLVs (Figures 2D & F), demonstrated dramatic recovery of these outcome measures to WT levels when treated with L-NAME (Figures 4D & F) indicating a role for nitric oxide in these defects. Finally, L-NAME treatment dramatically improved the contraction frequency of both Expanding and Collapsed PLVs beyond WT levels, but this was significant only at the highest two pressures (8-10 cmH_2_O).

**Figure 4.**
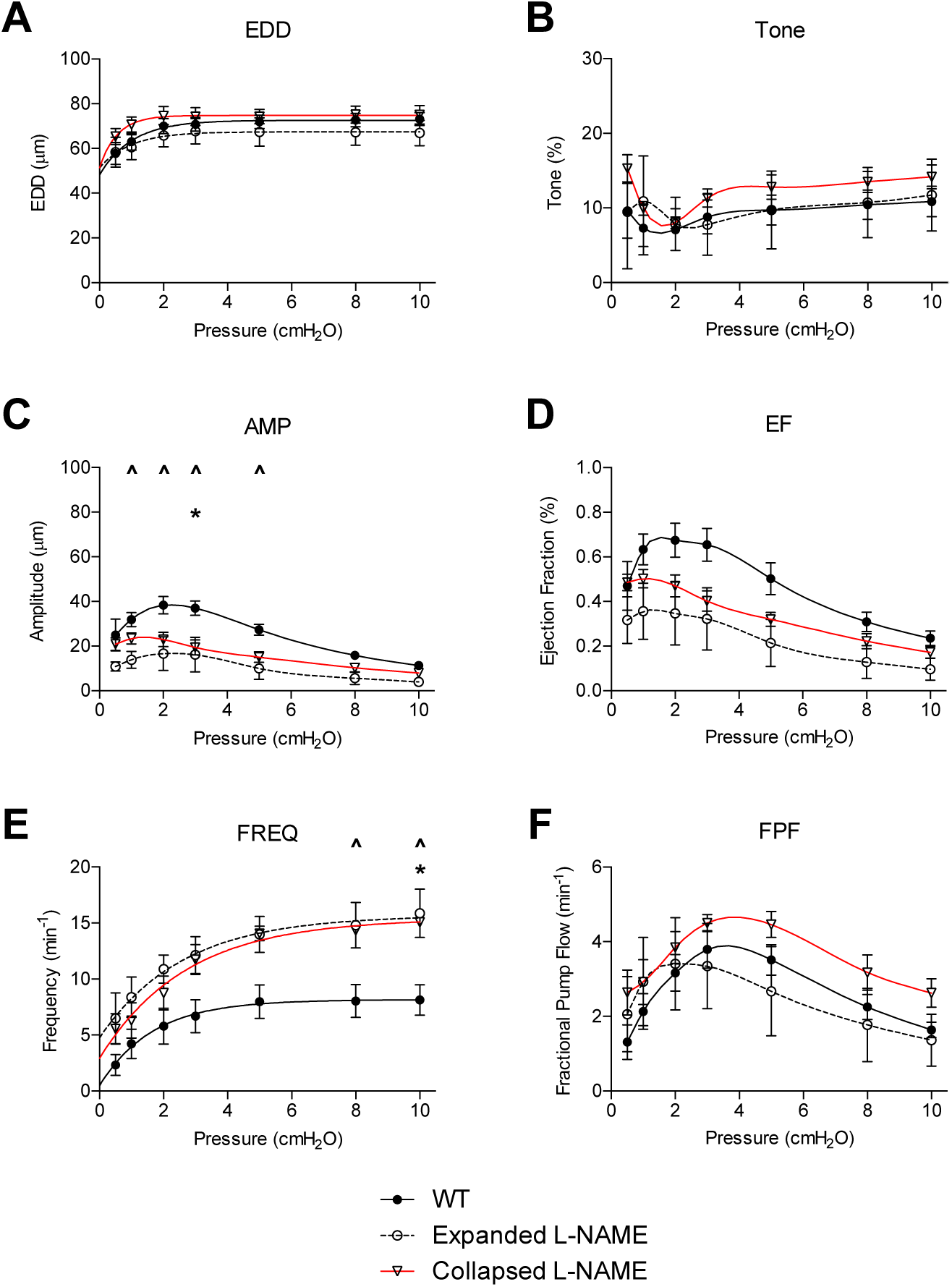
Nitric oxide synthase inhibition corrects TNF-Tg lymphatic vessel defects in contractile function. The lymphatic vessels described in Figure 1 were treated with L-NAME (10^−4^ M) and assessed for EDD, % tone, AMP, EF, FREQ, and FPF at the indicated pressures. All data are means (± SEM). When error bars appear missing, they are contained within the data points (\**P* < 0.05 Collapsed+L-NAME vs. WT; ^*P* < 0.05 Expanded+L-NAME vs. WT).

## DISCUSSION

Many RA patients suffer from periodic arthritic flares, but the mechanisms that promote and sustain bouts of joint inflammation remain poorly understood. Recent investigations suggest a contribution from circulating IgD^+^ immature B lymphocytes and PRIME cells, as well as synovial macrophage populations (8, 9). These novel observations are also consistent with our model of lymphatic dysfunction in RA flare (7), as: 1) the circulating IgD^+^ B-cells may be the B-in cells that we observe within the lymphatic sinuses (20, 21, 35), 2) the circulating PDPN^+^ PRIME cells may be lymphatic endothelial progenitors that are mobilized to repair damaged lymphatic vessels, and 3) the activated adherent macrophages in the lumen of degenerated PLVs (42) are likely from the afferent inflamed synovium (35, 43).

We have focused on the synovial lymphatic system during arthritic progression and flare (10, 12, 16, 20, 25, 26, 35, 43, 44), and the effects of interventions that specifically target lymphatic contractions, whose physiologic importance in supporting lymphatic drainage and subsequent onset of lymphedema have been established in preclinical and clinical studies (7, 29, 45, 46). We have also performed transmission electron microscopy studies on WT, Expanding and Collapsed PLVs (42). These studies confirmed that large, activated macrophages attach to damaged endothelial cells in Expanding PLVs of TNF-Tg mice with early arthritis, and that lymphatic muscle cells undergo apoptosis in Collapsed PLVs of TNF-Tg mice with advanced arthritis. Remarkably, this TNF-mediated damage was reversible with anti-TNF therapy that ameliorated the erosive inflammatory arthritis, in part via restoration of PLV contractions and potential enhancement of inflammatory cell egress (42). However, while these studies strongly implicate intrinsic PLV defects in the progression and flare of joint disease in TNF-Tg mice, in vivo studies cannot isolate PLV function independent of the upstream inflammation in the arthritic joint, and potential immune reactions in the efferent PLN.

Here we utilized ex vivo methodologies to directly assess functional differences between PLVs from TNF-Tg mice with Expanding and Collapsed PLNs, versus PLVs from their WT littermates. The results indicated significant deficiencies in TNF-Tg PLVs, which formally demonstrates the intrinsic defects in joint draining lymphatic function predicted by in vivo imaging and histological analyses. Thus, the lack of ICG clearance following foot pad injection of mice with inflammatory arthritis (7), and in the hands of patients with symptomatic RA (29), is likely due to the loss of lymphatic vessel contractile function that we observed as decreases in AMP, EF, and FPF. In addition, non-selective inhibition of NOS using L-NAME demonstrated recovery of AMP, EF, and FPF in the Collapsed TNF-Tg PLVs, but the vessels continued to exhibit decreased AMP during the Expanding phase. The specific contributions of endothelial nitric oxide (eNOS) and iNOS on these outcome measures through administration of L-NIL would be interesting to investigate in future studies given the proposed isolated role of iNOS on the deficits of TNF-Tg lymphatic contractility in vivo (25). Of note, the retained deficiency in AMP during the Expanded phase following NOS inhibition ex vivo (Figure 4C), and the incomplete resolution of lymphatic function following global iNOS ablation in vivo (25), suggests additional mechanisms are likely associated with the deficiencies in PLV contractility during the progression of inflammatory arthritis (i.e. structural damage and lymphatic muscle cell apoptosis (42)), which is an active area of investigation.

We also observed some apparent inconsistencies with the in vivo data, most notably the absence of functional differences between Expanding and Collapsed PLVs, and the normal contraction frequency of Collapsed PLVs ex vivo. To interpret these discrepancies correctly, it is critical to note that Collapsed PLVs in vivo are filled with static monocytes and macrophages (35, 42), which are removed during the cannulation and preparations prior to ex vivo analysis. Thus, it may be that these resident immune cells, known to express high levels of iNOS and inflammatory cytokines (25), may be responsible for additional or more dramatic PLV deficits that were not detected in our ex vivo study. In addition, contraction frequency measured in vivo by NIR-ICG imaging requires lymphatic contractions with sufficient force to generate bolus flow of ICG through the vessel. While the pacemaker activity seems to remain intact in Collapsed PLVs based on the unchanged contraction frequency ex vivo (Figures 1C & 2E), other factors, such as a proposed lymphatic valvular insufficiency in Collapsed PLVs (11, 35), could not be assessed in our ex vivo studies. Another potential confounding caveat is that the additional environmental or functional factors of the PLVs in vivo, coupled with the reduced AMP, EF, and FPF noted ex vivo, may not produce sufficient fluid flow to be visualized by in vivo NIR-ICG imaging. Thus, while the measures of in vivo and ex vivo contraction frequency may seem inconsistent for Collapsed PLVs, the in vivo measures may represent a loss of “effective” contractions associated with successful anterograde lymphatic drainage.

In summary, we show for the first time that PLVs from TNF-Tg mice with inflammatory arthritis have intrinsic functional defects that result in significant contractile dysfunction. As these defects have been associated with arthritic progression and flare in animal models of RA and patients with symptomatic disease, these data provide direct evidence of isolated lymphatic vessel dysfunction, and further validate lymphatics as a therapeutic target. Thus, drugs and stem cell therapies specifically designed to improve lymphatic vessel repair and the return of homeostatic lymphatic contractions are potential treatments for RA that warrant future investigation.

## ACKNOLWEDGEMENTS

This work was supported by research grants from the National Institutes of Health (T32s GM007356 and AR076950; R01s HL-12256, HL-120867, HL-142905, AR069000 and AR056702; and P30 AR069655).

